# Recombination rate and efficiency of linked selection in small and large stickleback populations

**DOI:** 10.64898/2026.03.18.712813

**Authors:** Hongbo Wang, Chaowei Zhang, Kerry Reid, Juha Merilä

## Abstract

Population genetic theory predicts that natural selection will be more efficient in large than small populations because genetic drift reduces the efficiency of selection in small populations. Small populations adapting to new environments can also be expected to evolve higher recombination rates to facilitate adaptation as well as to dissociate and purge harmful mutations. We tested these hypotheses (1) by investigating differences in the strength of association between nucleotide diversity (*π*) and recombination rate across the genomes of nine-spined sticklebacks (*Pungitius pungitius*) from four small freshwater (mean *N_e_* ≈ 2 578) and four large marine (mean *N_e_* = 86 742) populations, as well as (2) by comparing recombination rates between small and large populations using population specific linkage maps. We found the predicted positive correlation of *π* with recombination rate from all but the smallest freshwater populations, suggesting prevalent linked selection even after accounting for variation in GC/CpG content, and gene density. Mean recombination rates did not differ between freshwater and marine populations, except that the smallest *N_e_* freshwater population exhibited significantly elevated recombination rate. GWAS analyses suggested a polygenic basis for recombination rates. These results suggest an important role for linked selection in reducing *π* in low recombination regions especially in large populations. Moreover, as predicted by theory, at least one of the small freshwater populations appears to have evolved a higher recombination rate than its marine ancestors.

## Introduction

Genomic patterns of neutral variation are influenced by the forces of genetic drift, inbreeding, mutation and selection. Both positive selection favouring beneficial mutations (selective sweep) and negative selection against deleterious mutations (background selection) can reduce genetic diversity at genomic regions linked to targets of selection (Maynard Smith and Haigh 1974; Hudson and Kaplan 1995; McVean and Charlesworth 2000; Comeron et al. 2008; Becher and Charlesworth 2025). Given that reductions in genetic diversity are expected to be the most pronounced when selected and neutral mutations are tightly linked, one would expect to see a positive correlation between nucleotide diversity and recombination rate across the genome (Kaplan et al. 1989, Hudson and Kaplan 1995). Although this expectation has been verified in several studies of mostly well-established model organisms (e.g. Begun and Aquado 1992; Tenaillon et al. 2002; Cai et al. 2009; Geraldes et al. 2011; Corbett-Detig et al. 2015), the correlation is not universal (Cutter and Payseur 2013; Kartje et al. 2020) and studies of natural populations of non-model organisms are still rare. Therefore, as pointed out by Kartje et al. (2020) there is a need to evaluate this relationship in additional species and populations to better understand the causes of conflicting patterns across the studies. In particular, given that the efficiency of selection is expected to be greatly reduced in small populations (e.g. Petit and Barbadilla 2009; Charlesworth 2009), it would be of particular interest to test whether the diversity-recombination correlation is less pronounced for small as compared to large populations of the same species.

Recombination rates are heritable (Kivikoski et al. 2023a) and are known to evolve (Smukowski and Noor 2011; Dapper and Payseur 2017). There are also theories predicting that small populations adapting to new environments can be expected to evolve higher recombination rates (Felsenstein and Yokoyama 1976; Otto and Barton 2001). This is because an increased recombination can generate novel combinations of alleles or traits (Burt and Bell 1987) and allow for easier dissociation and purging of deleterious alleles. In particular, when multiple linked loci linked to recombination rate modifiers are repeatedly under moderate to strong selection, an increased recombination can evolve (Otto and Barton 2001; Bursell et al. 2025). However, this increase is expected only under restrictive conditions involving small population sizes, strong and consistent directional selection, high beneficial mutation rates, cis-acting recombination modifiers, and minimal recombination between selected loci (reviewed in Bursell et al. 2025). Whether these conditions are met in the wild is not generally known, and empirical evidence for adaptive divergence in recombination rates is scarce.

While there are several studies which have compared recombination rates among different populations of the same species (e.g. Shanfelter et al. 2019; Samuk et al. 2020; Venu et al. 2024), or between different subspecies (Dumont et al. 2010), these have all been two-population comparisons, limiting the inference that can be drawn from environmental drivers of observed differentiation or lack thereof. Similarly, some of these studies have used linkage disequilibrium-based methods subject to potential confounding effects of demography and selection. To the best of our knowledge, the strongest evidence for adaptive differentiation in recombination rates comes from the study by Samuk et al. (2020), who found an 8% difference in genome-wide recombination rates between two US populations of *Drosophila pseudoobscura*. Given all this, there is a call for studies including multiple populations, utilizing linkage maps estimated from family crosses (to avoid biases due to demography and selection; Raynaud et al. 2023) to compare recombination rates among populations of varying effective population sizes.

The aims of this study were two-fold. First, to evaluate the predicted positive correlation between nucleotide diversity (*π*) and recombination rate, and in particular, whether the strength of this relationship differs among populations with small and large effective population sizes (*N_e_*). Second, to compare the recombination rate between these small and large populations to test the prediction that small populations would have evolved higher recombination rates than larger populations. These predictions were tested in the nine-spined stickleback (*Pungitius pungitius*) using data from four natural populations with small *N_e_* and four natural populations with large *N_e_.* As the small *N_e_* populations of this species have been subject to recent strong directional selection (e.g. Herczeg et al. 2009, 2010; Karhunen et al. 2014; Kemppainen et al. 2021; Fraimout et al. 2022; Yi et al. 2024) enhancing the possibility of evolution of high recombination rates (Bursell et al. 2025). To enhance the accuracy genome-wide nucleotide diversity and recombination rate estimates, we utilised a high-quality gapless telomere-to-telomere reference genome (Zhang et al. 2026).

## Results

### Linkage maps and female-biased heterochiasmy

In this study, we investigated recombination rates and landscapes of *P. pungitius* from four small freshwater and four large marine populations of *P. pungitius* (Figure 1A; Table 1). A total of ten high-density linkage maps were generated for the two ecotypes and eight populations based on 158,980-3,838,162 SNP markers (Table 2). While the marker density in freshwater populations was lower than that in marine populations (t-test, t_105.3_=-31.67, p<2e-16; Table 1), the genomic range covered by these markers (i.e. the physical distance) was similar between the two ecotypes (t-test, t_158_=-0.077, p=0.94; Table 1), covering more than 99% genomic area of the autosomal regions (426.65-427.89 Mb out of the total 429.33Mb). The estimated genetic map lengths ranged from 1,556 to 1,815 cM for autosomes (Supplementary Figure 1-2; Figure 1; Table 2). There was a significant positive correlation between sex-averaged genetic map length (cM) and physical distance (Mb) at the chromosome level when pooling data from all populations (r_s_=0.61, df =158, p=2.03e-16; See Supplementary Figure 3A for the separate correlations of the 10 linkage maps).

**Figure 1.**
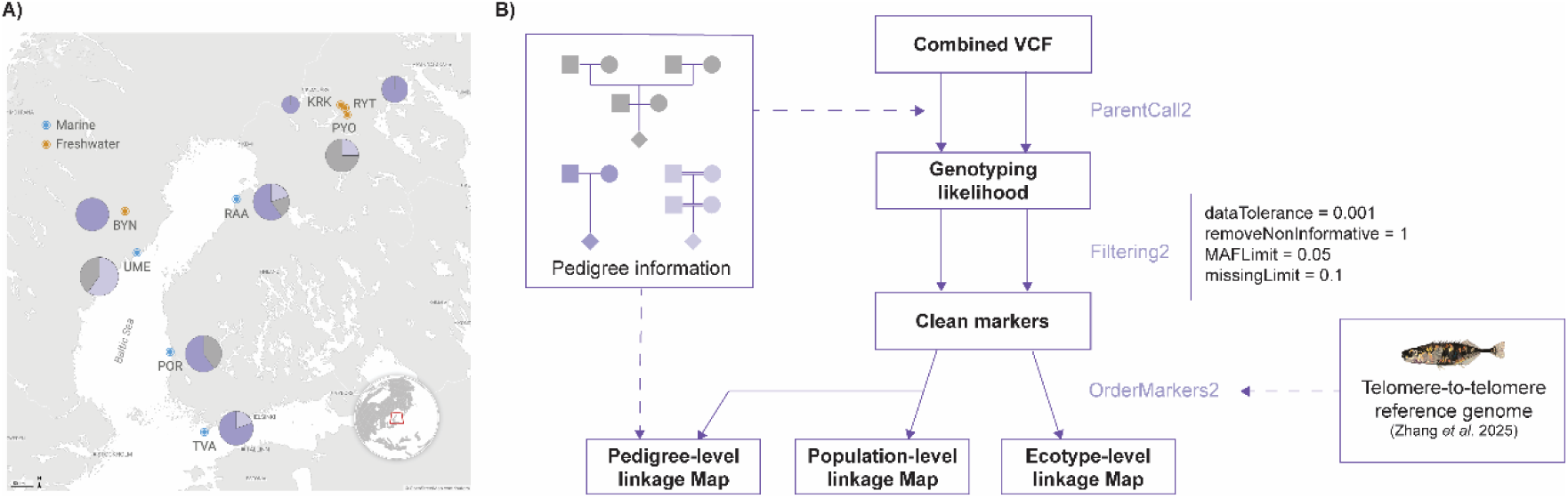
A) A map of the sampling sites of this study. The pie charts indicate the compositions of pedigree types (the different colour in pie diagrams depict the three pedigree types shown in B), and the circle sizes depict the number of individuals analysed in each population. Detailed information please refer to Supplementary Table 3. B) A summary diagram of the pipeline applied to construct the linkage maps at the ecotype and population levels.

**Table 1.** Basic information on nine-spined stickleback populations and samples used in this study. Ne_MSMC2-hm_=the harmonic mean of effective population size estimated from 2 - 200 kya by MSMC2. Ne_CurrentNe2_= effective population size estimated by CurrentNe2. *π* = mean nucleotide diversity estimated from wild-caught parents. n_family_ = number of families, n_F0_: number of wild caught parental fish, n_offspring_: number of F1 or F2 offspring. Ecotype MA: Marine, FW: freshwater.

| Population | Code | Coordinate | Ecotype | $N_e$-<br>MSMC2-hm | $N_{e-CurrentNe2}$ | $\pi$ | $n_{family}$ | $n_{F0}$ | $n_{offspring}$ |
| --- | --- | --- | --- | --- | --- | --- | --- | --- | --- |
| Pori | POR | 61.59111°N,<br>21.47295°E | MA | 261179 | 12972.6 | 0.0049 | 5 | 12 | 54 |
| Raahe | RAA | 64.68818°N,<br>24.46189°E | MA | 348498 | 10925.9 | 0.0051 | 5 | 12 | 54 |
| Tvärminne | TVA | 59.83333°N,<br>23.0°E | MA | 255481 | 24262.0 | 0.0050 | 5 | 10 | 52 |
| Umeå | UME | 63.63739°N,<br>19.99438°E | MA | 194543 | 9136.8 | 0.0050 | 5 | 10 | 58 |
| Bynastjärnen | BYN | 64.45416°N,<br>19.44075°E | FW | 8648 | 403.9 | 0.0018 | 5 | 10 | 33 |
| Kirkasvetinenlampi | KRK | 66.43673°N,<br>29.13568°E | FW | 11389 | 266.5 | 0.0014 | 4 | 8 | 24 |
| Pyöreälampi | PYO | 66.26226°N,<br>29.42916°E | FW | 4235 | 18.4 | 0.0011 | 5 | 10 | 49 |
| Rytilampi | RYT | 66.38482°N,<br>29.31561°E | FW | 9784 | 384.0 | 0.0019 | 4 | 8 | 40 |

**Table 2.** Details of the ten linkage maps. Marker number= number of autosomal markers applied after filtering, Physical distance = the valid physical distance covered by markers (Mb), Map length = the estimated genetic distance (cM), Recombination rate = the genomewide recombination rate (cM/Mb) with confidence intervals calculated from chromosome-level estimates, Heterochiasmy ratio = female-to-male ratio of the recombination rate. FW-all and MA-all refer to the freshwater and the marine linkage maps, respectively.

| Linkage map | Marker number | Physical distance (Mb) | Map length (cM) | Recombination rate (cM/Mb) | Heterochiasmy ratio |
| --- | --- | --- | --- | --- | --- |
| PYO | 158,980 | 426.65 | 1,815 | 4.24 (4.01 - 4.68) | 1.59 |
| KRK | 578,795 | 426.26 | 1,664 | 3.90 (3.71 - 4.14) | 2.08 |
| RYT | 884,701 | 427.18 | 1,584 | 3.71 (3.48 - 4.04) | 2.12 |
| BYN | 877,575 | 426.88 | 1,605 | 3.76 (3.56 - 4.03) | 1.79 |
| TVA* | 3,838,162 | 427.89 | 1,633 | 3.82 (3.63 - 4.10) | 2.02 |
| RAA | 3,813,481 | 427.73 | 1,557 | 3.64 (3.47 - 3.84) | 1.95 |
| UME | 3,485,604 | 427.86 | 1,576 | 3.68 (3.53 - 3.89) | 1.58 |
| POR | 3,802,726 | 427.79 | 1,632 | 3.81 (3.66 - 4.06) | 2.14 |
| FW-noPYO** | 996,472 | 427.52 | 1,591 | 3.72(3.57 - 3.96) | 1.99 |
| FW-all | 1,041,122 | 427.55 | 1,743 | 4.08 (3.91 - 4.38) | 1.73 |
| MA-all | 1,784,241 | 427.74 | 1,556 | 3.64 (3.53 - 3.81) | 2.03 |

\*The population of origin for the reference genome.
\*\*FW-noPYO refers to the combined freshwater linkage map excluding the PYO population

Female genetic map lengths were longer than those of the males at the chromosomal levels (paired t-test, t_199_=31.16, p=2.2e-16), with an average female-to-male map ratio of 1.73 in freshwater (FW) and 2.03 in marine (MA) linkage maps, respectively. This heterochiasmy was present in all chromosomes (Supplementary Figure 2). The most differentiated chromosome in terms of map length between the two sexes was the X chromosome (chrX). The maps for SDR consisted of 2,136 (FW) and 6,953 (MA) informative markers, whereas those of the pseudoautosomal region (PAR) had 94,252 (FW) and 296,375 (MA) markers.

While no recombination should be observed on male SDR, the genetic distances were estimated to be 106.5 (FW) and 119.7 cM (MA) in female SDRs. Female map lengths exceeded those of males also in PAR (FW: 2.00 and MA: 1.63 times longer).

### Genomewide recombination rate variation among populations

The number of crossovers identified in individual offspring varied from 18 to 52, averaging at 21.1 maternal and 11.2 paternal crossovers per meiosis. The mean crossover number was significantly higher in freshwater than in marine populations (t-test, *t*_252_= −5.27, p = 2.94e-07; Figure 2A, Table 2), and PYO exhibited the highest crossover count (x = 40.30; CI: 38.90–41.70) that is significantly higher than the other populations (t-test, t_947.16_ = −8.97, p < 2e-16, Supplementary Figure 12A). It demonstrated 1.36 times more crossovers than in UME, which had the lowest number among the eight populations (x = 29.66; CI:28.22–31.10; Figure 2B). The population-level number of crossovers was negatively correlated with the past *N_e_* estimated by MSMC2 (generalised linear mixed model [GLMM], β = −1.58, p = 0.0022), but not significantly so with contemporary *N_e_* estimated by CurrentNe2 (GLMM, β = −0.92, p = 0.098; Figure 2C). On average, 1.84 crossovers were detected on each autosomal chromosome per meiosis, and the crossover numbers were a positive function of chromosome length, and the association was shallower in males than in females (GLMM, β_chr-length_ = 0.18, p_chr-length_ < 2e-16; β_chr-length:sex_ = −0.14, p_chr-length:sex_ < 2e-16, Supplementary Figure 3B). The chromosomal recombination rates ranged from 1.22 to 7.18 cM/Mb, and they scaled negatively with chromosome length (GLMM, β = −0.24, p = 1.69e-11; Figure 2D).

**Figure 2.**
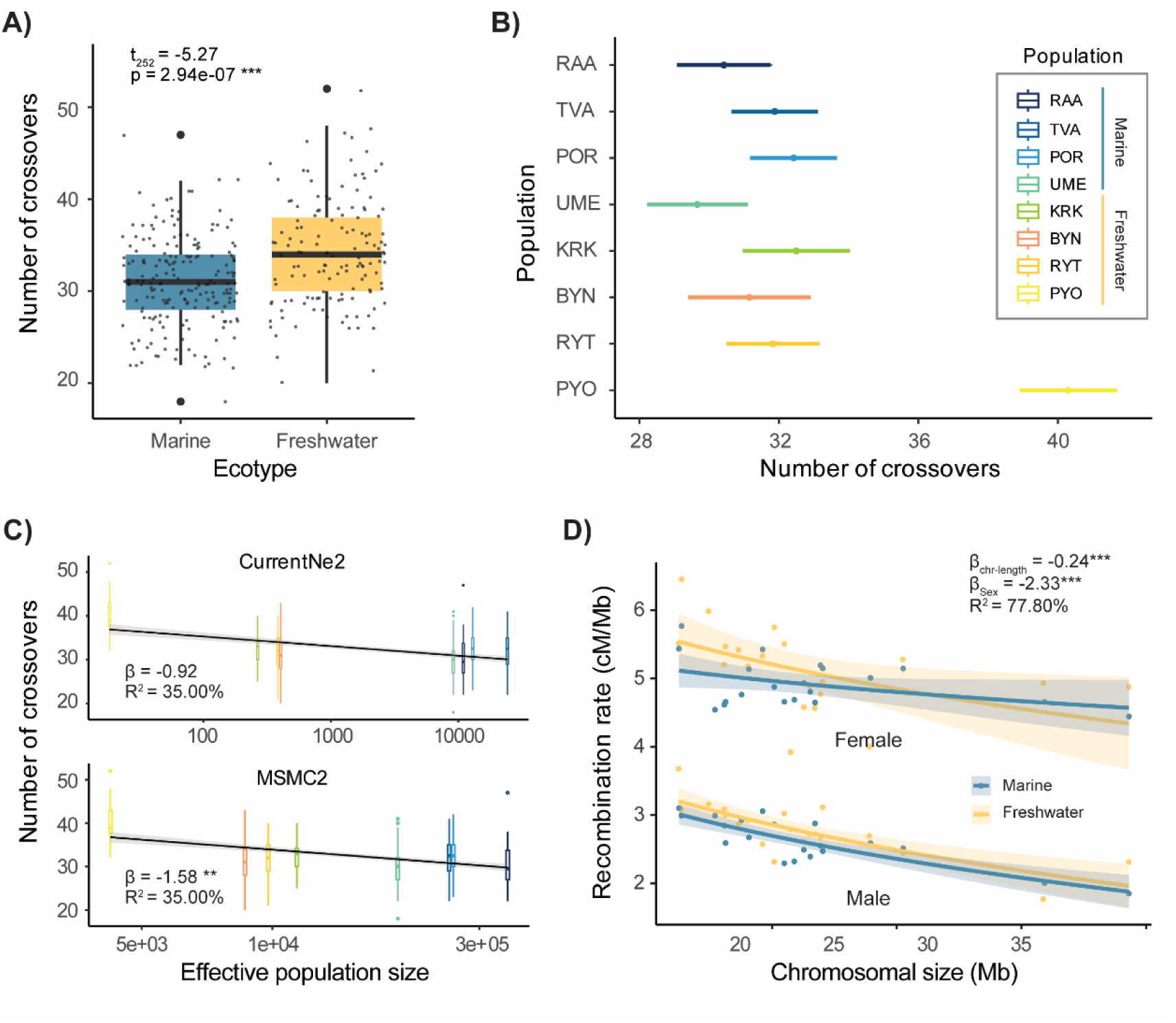
A) Mean crossover number in the marine and freshwater samples (p-values derived from t-test). B) Genomewide recombination rates for the eight population-level linkage maps, with the error bars depicting 95% confidence intervals based on chromosome-level estimates. C) Negative correlation between crossover number and the effective population size (*N_e_*); upper panel = *N_e_* estimated from CurrentNe2, lower panel = *N_e_* estimated from MSMC2. D) Male and female recombination rates as functions of chromosome size in marine and freshwater populations. The estimated slopes (β) and variance explained [R^2^] are shown for panel C and D, which were derived from the generalised linear model for panel C and the optimal generalised linear mixed model with population as the random effect for panel D using glmmTMB. Significance levels: <0.001: ***, <0.01: **, < 0.05: *.

The sex-averaged recombination rates estimated from freshwater and marine linkage maps did not differ significantly from each other (MA = 3.74, FW = 3.91 cM/Mb; t-test, t_3.93_=1.24, p=0.28; Table 2). Freshwater populations exhibited slightly higher chromosomal recombination rates compared to the marine populations (MA = 3.63 [95% CI 3.53 - 3.81], FW = 4.07 [95% CI 3.91 - 4.38] cM/Mb; t-test, t_141.69_ = 2.14, p = 0.03), although this difference was no longer significant upon removal of PYO individuals (FW_non-PYO_ = 3.72 cM/Mb, t-test, t_113.06_=-0.61, p=0.55, Supplementary Figure 12B).

### Localised recombination rates are conserved

Despite the differences in the overall recombination rates, the genetic maps varied similarly throughout the genome for the eight populations (Supplementary Figure 1), with the 1Mb-window recombination rates tightly aligned between the two ecotypes (r = 0.74, df = 903, p < 2.2e-16) and the two sexes (r = 0.54, df = 904, p < 2.2e-16; Supplementary Figure 7). In general, recombination ‘hotspots’ tend to be located at the telomeric regions of the chromosomes (Figure 3A; Supplementary Figures 4 and 8). These regions are typically characterized by high methylation levels. The CpG content was significantly higher in the identified recombination ‘hotspots’ compared to ‘coldspots’ (t_40.93_= −14.39, p < 2.2e-16; Figure 3A, Supplementary Figure 8). Recombination rates were strongly and positively associated with CpG levels in the 1Mb windows (negative binomial GLMM, Figure 4A) with a significant sex-specific effect of methylation level on recombination rate (negative binomial glmmTMB model, Fig 4A). Conversely, regions with high gene density exhibited fewer recombination events, indicating a weak negative association between gene density and recombination activity (negative binomial GLMM, Figure 4B). A specific analysis of the well-characterised inversion region on chr19 (Varadharajan et al. 2019; Kivikoski et al. 2021) also revealed that the structural variations suppressed crossovers (Supplementary Figure 9).

**Figure 3.**
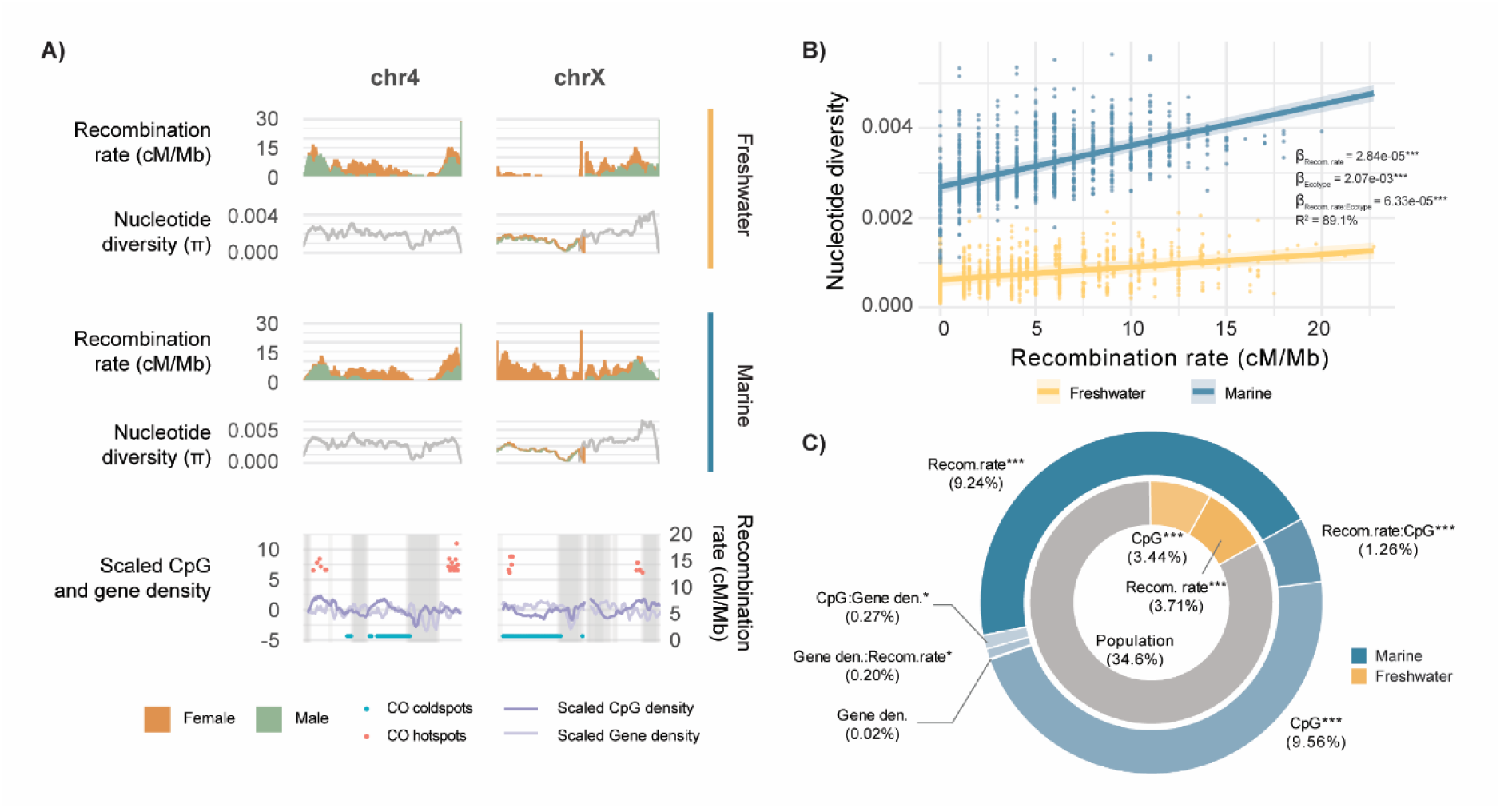
A) Localized patterns of recombination rates (cM/Mb) and nucleotide diversity (π) in the two ecotypes as exemplified by chromosome 4 and chromosome X across 1Mb sliding windows (orange: females, green: males; see Supplementary Figure 5). The bottom panel displays scaled CpG density and scaled gene density, with recombination hotspots and coldspots marked, and repeat regions shaded in gray (also see Supplementary Figure 8). B) Association between nucleotide diversity (π) and recombination rate (cM/Mb) for the two ecotypes. β refers to the estimated slope coefficient for each variable incorporated in the optimal model. Significance levels: <0.001: ***, <0.01: **, < 0.05: *. C) The proportion of variance in 1 Mb window-sized nucleotide diversity (π) explained by the variables in the optimal linear model for freshwater (outer) and marine (inner) populations, including recombination rate (cM/Mb), CpG content, their interactive effect, and the random effects.

**Figure 4.**
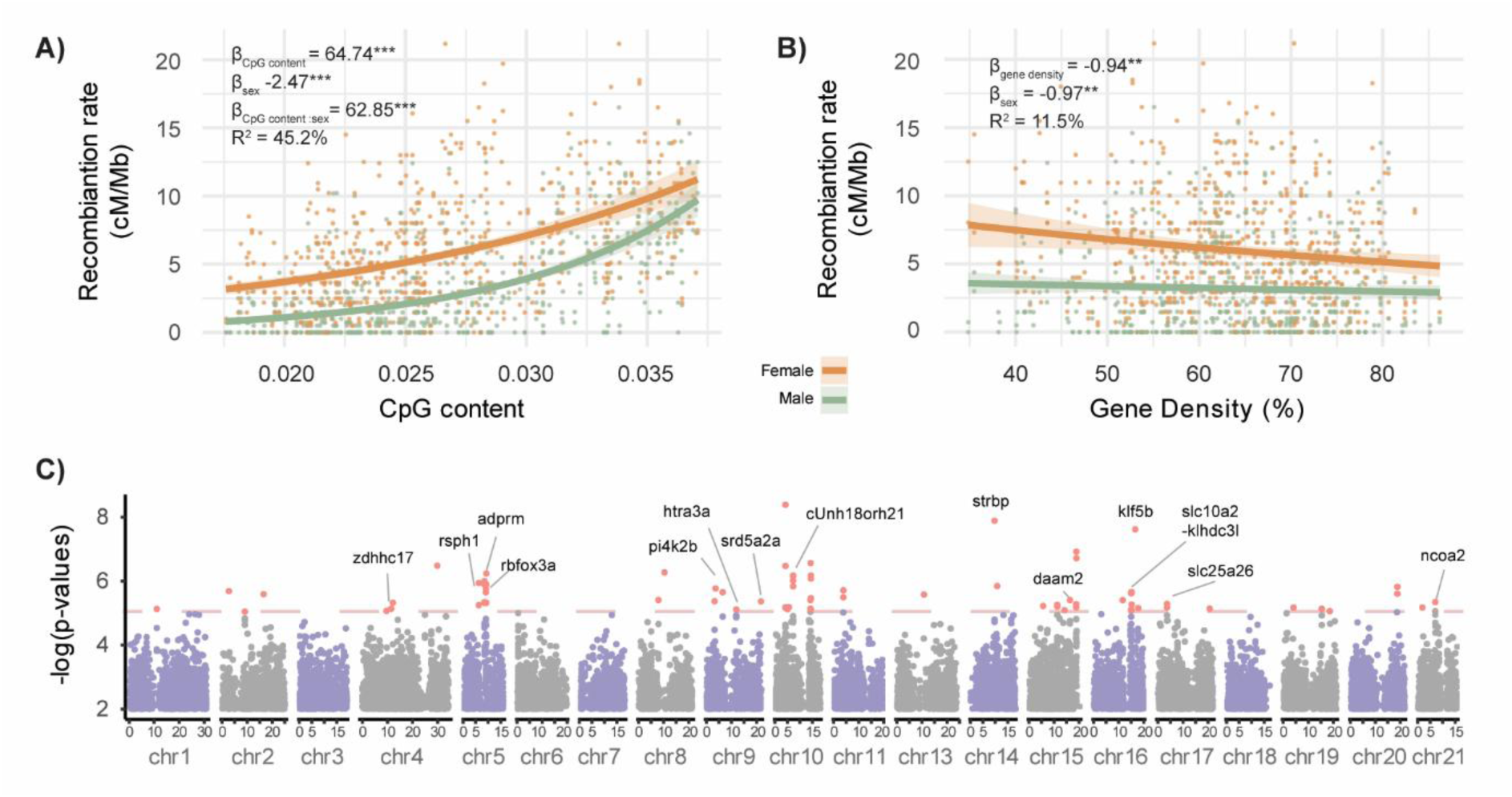
Localised recombination rates (cM/Mb; in 1 Mb windows) associated with A) CpG content and B) gene density. Statistical analyses were conducted using negative binomial GLMMs implemented in glmmTMB. β denotes the estimated slope coefficient for each variable incorporated in the optimal model. Significance levels: <0.001: ***, <0.01: **, < 0.05: *. C) The Manhattan plot for genome wide association analysis conducted with FarmCPU, where the red dots above the dotted line are the SNP sites with p-values < 1e-05. The available gene annotations are marked around those sites.

### Nucleotide diversity (π) and recombination rate

The optimal generalised linear model (GLM) models explaining variation in localised π, based on population- and ecotype-specific linkage maps, explained 88.77% and 53.52% of the variance in π across 1Mb non-overlapping windows, respectively. Because the average *π* was higher in marine than in freshwater populations (Table 1), ecotype accounted for the majority of variance in both models – 83.68% in the population-level linkage maps and 43.63% in the ecotype-level linkage maps (Table 3). Nevertheless, both models indicated that the localised *π* was positively associated with sex-averaged recombination rate and CpG content, while it was weakly negatively associated with gene density (Table 3). These findings were consistent with simpler models incorporating only one of these genomic features alongside Ecotype (Figure 3B, Supplementary Figure 11). Furthermore, this positive association between *π* and recombination rates was significantly shallower in freshwater than in marine populations (GLMM, β_recom.rate:ecotype_ = 6.33e-05, p < 2e-16, Figure 3B), and interestingly, the most inbred population in this study, PYO, was the only population that did not exhibit a significant association between the two (β = 2.91e-06, p=0.43, Supplementary Figure 5).

**Table 3.** A summary of the best multivariate models for the factors influencing localised nucleotide diversity (RR = sex-averaged recombination rate). All explanatory factors are standardised using Z-score normalisation. (Significance levels: <0.001: ***, <0.01: **, < 0.05: *)

|  |  | Population-level |  | Ecotype-level |  |
| --- | --- | --- | --- | --- | --- |
| | | Slope<br>($\beta$, significance<br>level) | Variance<br>explained | Slope<br>($\beta$, significance<br>level) | Variance<br>explained |
| Fixed main effect | Ecotype | 1.82*** | 83.68% | 1.31*** | 43.43% |
|  | RR | 0.07*** | 0.98% | 0.25*** | 6.52% |
|  | CpG content | 0.06*** | 1.04% | 0.20*** | 2.32% |
|  | Gene density | -0.006 | 0.01% |  |  |
| Fixed interaction<br>terms | CpG:recom. | -0.03*** | 0.06% | -0.07** | 0.38% |
|  | Ecotype:RR | 0.12*** | 0.83% | 0.19*** | 0.87% |
|  | CpG:ecotype | 0.12*** | 0.86% |  |  |
|  | Gene density: RR | -0.02* | 0.00% |  |  |
|  | CpG:gene density | 0.02** | 0.02% |  |  |
| Random effect | Population |  | 1.29% |  |  |
| Intercept |  | -0.89*** |  | -0.63*** |  |

To further examine other factors affecting localised *π*, we divided the data according to the ecotype. In this analysis, the models of both ecotypes revealed that recombination rate scaled positively with *π* and together with CpG contents explained the most variance among all fixed effects (Figure 3C). However, the freshwater populations exhibited much greater residual variation (FW: 34.6%, MA: 0.0%; Figure 3C), indicating that *π* is more variable among freshwater than in marine populations, weakening the association between *π* and recombination rate.

### Genetic loci associated with crossover numbers

Although there were differences in the ability of different approaches to identify crossover-associated loci, approximately 4 to 48 putative quantitative trait loci (QTL, SNPs within 100kb from each other consider to be same QTL) were found to be associated with genomewide crossover numbers (a nominal significance threshold p < 1e-05), suggesting a polygenic basis for recombination rate variation. In MLM and GLM, 4 and 48 QTL were identified to be associated with recombination rates with a threshold of p < 1e-05, respectively. The FarmCPU module, which effectively controls for false positives and maintains high detection power (Liu et al. 2016), identified 48 suggestive QTL (p < 1e-05, Figure 4C) distributed across the genome (Figure 4C) involving 26 genes with unknown functions, but also several well-studied candidate genes such as *adprm*, *ncoa2*, *htra3a*, *mbtps1* and *slc25a26*. Six QTL were located in the intergenic regions, including those within *slc10a2-klhdc3l*, *rsph1-adprm*, and rbfox3a-LOC134133006 (uncharacterised). To rigorously account for multiple testing while maintaining statistical power, we applied the Benjamini-Hochberg False Discovery Rate (FDR) correction. At a threshold of FDR < 0.05, three QTL remained significantly associated in both the FarmCPU and GLM analyses, whereas none remained significant in the MLM.

## Discussion

The most salient findings of this study include the consistent and strong positive association between nucleotide diversity and recombination rate in large outbred marine populations, as well as the significantly shallower positive association of these parameters in small inbred freshwater populations. Likewise, the higher crossover counts and recombination rates in the smallest freshwater population compared to that in large marine populations suggest that it may have evolved higher recombination rates. The results further confirm a clear female-biased sex heterochiasmy in crossover counts, and that the crossovers are more prevalent in the chromosome ends, in the regions with high CpG content and low gene density. We discuss these findings and their implications to our understanding of genomewide and localised patterns of genetic diversity and evolution of recombination rate variation.

The observed positive correlation between nucleotide diversity and recombination rate is expected if background selection or selective sweeps reduce genetic diversity in low recombination regions of the genome (i.e. Hill-Robertson interference; Hill and Robertson, 1966). This correlation was much weaker (or even absent, as in the case of the most homozygous population) in the small freshwater populations as a likely result of strong genetic drift prevailing in them. The effect of strong genetic drift is to reduce the efficiency of selection (Lynch et al. 2011) and thereby weaken the association between nucleotide diversity and recombination rate. This inference is supported by several lines of corroborative evidence. First, although pond populations have been able to purge much of their highly deleterious mutation load, they have accumulated relatively high loads of mildly deleterious mutations suggesting weakened efficiency of selection (Chen et al. 2025).

Second, pond populations are significantly enriched for young active transposable elements compared to marine populations (Varadharajan et al. 2019). Again, selection against repeats and elimination of accumulated repeat copies is expected to be reduced in small populations allowing active repeat families to expand rapidly in size (Lynch and Conery 2003). Third, the degree of population differentiation among the pond populations is very high (Shikano et al. 2010; Fraimout et al. 2022) as are their inbreeding levels (Kivikoski et al. 2023b; Chen et al. 2025). Furthermore, the current effective population sizes in ponds are more than a thousand-fold lower than in outbred marine populations (Feng et al. 2025; Zhang et al. 2026). All these observations suggest that the pond populations are subject to strong genetic drift, and that the weakened correlation between nucleotide diversity and recombination rate in ponds is likely a reflection of reduced efficiency of natural selection weakening the correlation between nucleotide diversity and recombination rate. All this said, it is also conceivable that population size bottlenecks experienced by freshwater populations could mask selection on linked sites (Beissinger et al. 2016; Lucena-Perez et al. 2021). However, there is also empirical evidence for the opposite - bottleneck amplifying signatures of selection (Torres et al. 2018).

The observation that the smallest pond population had higher crossover counts and recombination rate than the large marine populations may suggest that after colonising the freshwater habitats about 10 000 years ago (Feng et al. 2024), this pond population has evolved higher recombination rates. In fact, small populations adapting to new environments can be expected to evolve higher recombination rates as they would benefit from generation of novel combinations of alleles and traits (Otto and Barton 2001; Burt and Bell 1987).

Furthermore, as the pond populations suffer from elevated loads of deleterious mutations (Chen et al. 2025), increased recombination rates would facilitate easier dissociation and purging of deleterious alleles and the random linkage disequilibrium generated by genetic drift (Barton and Otto 2005; Charlesworth et al. 2009; Lynch et al. 2011). However, although the recombination rate was significantly higher in the most inbred freshwater population, the other freshwater populations did not exhibit elevated recombination rates. While this could be explained by weakened efficiency of selection in small *N_e_* freshwater populations constraining (polygenic) adaptation (Kimura 1963; Lanfear et al. 2014; Petit and Barbadilla 2009) and reducing the likelihood of adaptive transformations, one has to remember that all studied freshwater populations are highly inbred, the most inbred population (PYO), having 91% of its genome estimated as homozygous (Chen et al. 2025). This is noteworthy because the long runs of homozygosity can be expected to reduce the detection probability of recombination events (Rastas 2017). This suggests that we might have underestimated the recombination rates in freshwater populations. In the same vein, if the freshwater populations have smaller genome sizes than marine populations as our ongoing studies suggest (Wang et al, unpublished), the recombination rate estimates of freshwater populations based on physical distances might be downward biased as the used reference genome comes from a marine population.

Genome structure dictates the distribution of the recombination events. Despite the population- and sex- differences in genomewide recombination rates, the distribution of the crossovers along the genome appeared to be conservative occurring more often at the subtelomeric parts of the chromosomes than around the centromeres. This is likely due to crossover interference (CI), a phenomenon in which the occurrence of one crossover reduces the likelihood of another nearby crossover (Otto and Payseur 2019). CI tends to push crossovers toward the chromosomal ends, thereby leaving more room in the central regions for additional crossovers (Johnston 2024). This limits the maximum number of crossovers per chromosome leading to lower recombination rates in longer chromosomes as we also observed. Furthermore, structural variants, such as the inversion on chr19 in this study, tend to restrict the occurrence of the crossovers (Bartolomé et al. 2002; Rizzon et al. 2002). Further studies should look into inversion frequencies in the two ecotypes and see if the large marine populations carry more inversions than the smaller freshwater populations.

We found that recombination rate variation was in strong association with CpG content, CpG rich areas having elevated recombination rates. This was expected as the stickleback *Prdm9* has lost its functional domains KARB and SSXRD (Cavassim et al. 2022), a feature also confirmed in both freshwater (v8, ENA: GCA_902500615; Wang et al. 2024) and marine (NSPV9_T2T, NCBI: PRJNA1334546; Zhang et al. 2026) reference genomes of *P. pungitius*. Consequently, the significant influence of CpG content on fine-scale localised recombination rates was expected as recombination hotspots tend to predominantly concentrate in promoter-like regions, including CpG-rich areas (Cavassim et al. 2022).

Similar results are available also from other *prdm9*-independent species, such as canids (Axelsson et al. 2012) and birds (Singhal et al. 2015). The GWAS results further support the notion that *prdm9* does not control recombination rate variation in sticklebacks: recombination in sticklebacks appears to be a polygenic trait influenced by multiple quantitative trait loci with most candidate genes differing from those identified in mammals (Johnston 2024; McAuley et al. 2024). Two of the QTL, slc25a26 and klhdc3l are particularly interesting. The former is known to be directly involved in the methylation process and cell proliferation (Menga et al. 2017) whereas the latter is structurally similar to recombination activate gene 2 (rag2) in humans (NCBI gene description to geneID: 116138).

Finally, although we employed a high-quality, telomere-to-telomere reference genome and high-depth sequencing data to ensure accurate SNP calling (Zhang et al. 2025), regions with repetitive sequences remain challenging for precise variant detection using short-read data. Consequently, these regions were excluded from most of our analyses. Integration of long-read sequencing data would provide means to improve variant detection in these regions and gain even better estimates of nucleotide diversity and crossover rates. Such data could also provide insights on the roles of transposable elements and other structural variants in influencing recombination patterns and genetic diversity.

In conclusion, utilising high-quality linkage maps and an improved reference genome, we were able to gain insights on recombination rate variation and its genomic basis in eight stickleback populations with contrasting effective population sizes. Evidence was recovered for markedly reduced nucleotide diversity in low recombination regions of the genome suggesting an important role for linked selection in reducing diversity. The lower positive correlation between nucleotide diversity and recombination rate in small than large populations is suggestive of reduced efficiency of selection in small populations subject to strong drift. The highest recombination rate in the smallest *N_e_* population may suggest that it might have evolved elevated recombination rates possibly to counteract the negative consequences of reduced genetic diversity and elevated linkage disequilibrium. However, the possibility that recombination rate difference relates to genome size and structure differences cannot be excluded and warrant further study. The results further suggest that recombination rate variation in sticklebacks has a polygenic basis, but further studies are required to explore the detailed molecular mechanisms underlying the recombination variation and to identify the potential molecular drivers of the crossover distributions for species without *prdm9*.

## Methods

### Construction of Linkage maps

Families from eight different *P. pungitius* populations (four freshwater and four marine) were used in this study (Figure 1A, Table 1; see details about the sampling, DNA extraction, and SNP calling in Supplementary Materials and Methods). The per-sample gVCF files were combined and genotyped jointly for each population with the CombineGVCFs and GenotypeGVCFs modules in GATK (v4.3.0.0; Poplin et al. 2017). GLnexus (v1.4.1; Yun et al. 2021) was utilised to generate the ecotype-specific vcf files, for the marine and the freshwater populations.

The combined vcf files, at both the ecotype and population levels, were applied to construct the linkage maps following the standardised Lep-MAP3 pipeline (v 0.5; Rastas 2017; Figure 1B). Firstly, the genotyping likelihoods of the biallelic SNP markers in each vcf file were obtained using the ParentCall2 module (XLimit = 2). Secondly, a series of filters were applied (dataTolerance = 0.001, removeNonInformative = 1, MAFLimit = 0.05, missingLimit = 0.1) to retain the informative markers and to eliminate markers violating Mendelian expectation. Thirdly, the filtered markers were ordered in the OrderMarkers2 module with the Morgan function, based on their physical positions indicated by the reference genome. The genetic distance between markers was then calculated for each vcf file, utilising the provided pedigree information (Figure 1B). A special treatment was implemented to the sex determination region (SDR) of chrX (chrX:1–18.875Mb) by selecting only those markers that were informative from the maternal side (informativeMask=2), while the recombination rate in males was set to zero.

### Estimation of recombination rate and crossover count

The genome wide recombination rates (*r*, cM/Mb) were calculated by dividing the total genetic distance (cM) over the valid physical distance covered by the markers, and the localised recombination rates were calculated for 1Mb non-overlapping windows, employing a custom script available from: https://github.com/LeoHongboWANG/Recombination_in_Sticklebacks. This method was applied to all linkage maps constructed, for each population and ecotype, to estimate recombination rates at different levels. The recombination rate ‘hotspots’ were defined as the first 5% windows exhibiting the highest recombination rates, while the ‘coldspots’ were windows with no detected recombination. A special examination of the crossover density was performed for the earlier (Varandharjan et al. 2019) identified inversion on chr19 (Supplementary Materials and Methods). The total number of crossovers identified from each individual was reported along with the linkage maps in the log files. However, to conduct a more detailed examination of the crossover distribution, the pedigree-level linkage maps were constructed by providing only the pedigree information of a specific family in the third step of the previous section (Figure 1B). Variations in genetic distance along these maps indicated crossover events. The positions and the number of crossovers for each parent were then directly obtained by identifying the changes in genetic distance values in these pedigree-specific linkage maps.

### Nucleotide diversity, gene density and CpG contents

Population-level nucleotide diversity was estimated using SNPs from the F_0_ parents in 1Mb non-overlapping windows, executed with VCFtools (v 0.1.16; Danecek et al. 2011). Given that recombination rates are known to be associated with gene density and CpG sites (e.g. Axelsson et al. 2012), these parameters were estimated from the gene annotations and the methylation data of the T2T reference genome (Zhang et al. 2026) with the assistance of bedtools (v2.31.1, Quinlan et al. 2010) using 1Mb window size. Due to the possible inaccuracy of the SNP calling in the repeat regions, the genome regions with high repeat content (length proportion >20%) were excluded from these estimations.

### Statistical analyses

Linear mixed models were employed to assess the influence of effective population sizes (*N_e_*), estimated using MSMC2 and CurrentNe2 (see Zhang et al. 2026), on parental crossover count. After evaluation of the data distribution and appropriate scaling, the models were fitted using R package glmmTMB (v1.1.10; Brooks et al. 2017) with a ‘gaussian’ family with including pedigree as random effect. The rationale to analyse crossover counts rather than recombination rates was that our ongoing (and unpublished) research has revealed that freshwater populations have smaller genomes than marine populations. Since the used reference genome is from a marine population (Zhang et al. 2026), division with genetic map distance with marine genome size could bias the comparisons based on recombination rates. Therefore, we analysed the effect on effective population size (*N_e_*) on crossover counts using the model:

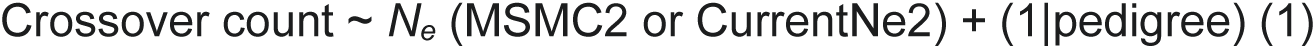

Similarly, mixed models were built to investigate the effect of chromosome length and per-chromosome crossover counts at population level. The initial model, incorporating parental sex and ecotype as covariates, was applied in the MuMIn R package (v1.48.4, Bartoń 2025), which evaluated all possible combinations of interaction terms and identified the best model based on the Akaike Information Criterion (AIC). The best model was:

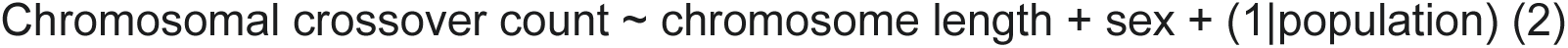

Furthermore, we explored pairwise associations between the localised nucleotide diversity (*π*), recombination rates (r, cM/Mb), gene density, and CpG contents. Following an examination of co-linearity of the explanatory variables in package ‘car’ (v3.1.1; Fox and Weisberg 2019), we investigated the impact of recombination rates on *π* through generalized linear mixed models (GLMMs) accounting for the influence of ecotype, CpG content and gene density. The aforementioned method was applied to determine the optimal model, while the variance explained by each term was assessed by dropping the given term and calculating the reduction of the explained variance from the original model – applying this method to either the main effect model (for the main effects) or the full model (for the interaction terms).

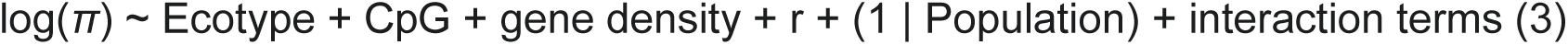

Because of the high impact of ecotype on π, we also constructed models separately to the two ecotypes.

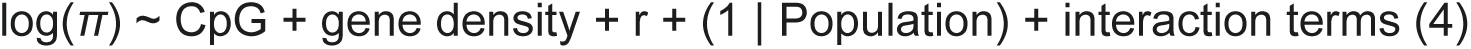

All best models were evaluated by plotting the residual and fitted values, while the dispersion estimates were examined to check for overdispersions.

### Genome wide association analyses

A genome-wide association study (GWAS) was carried out to assess the association between the parental genetic variants and the meiotic crossover counts. The analysis was conducted using the rMVP software (Yin et al. 2021), employing the General Linear Model (GLM), Mixed Linear Model (MLM), and Fixed and Random Model Circulating Probability Unification (FarmCPU) approaches, respectively. The first five principal components of the SNP matrix were included as covariates to exclude the impact of genetic relatedness in the models. Recombination rate-associated loci were identified using standard tests (p < 0.05) as FarmCPU is considered to be robust for handling population structure and effectively controlling for false positives caused by population stratification (Liu et al. 2016). To rigorously account for multiple testing, we applied the Benjamini-Hochberg False Discovery Rate (FDR) correction. We also note that FarmCPU is considered robust for handling population structure effectively controlling for false positives caused by population stratification (Liu et al 2016).

### Software availability

The custom scripts used for localized recombination rate calculations in this study are available on GitHub (https://github.com/LeoHongboWANG/Recombination_in_Sticklebacks).

## Data Access

All raw and processed sequencing data generated in this study have been submitted to the NCBI BioProject database (https://www.ncbi.nlm.nih.gov/bioproject/) under accession number PRJEB60682 and PRJNA1334546. The 10 sex-specific linkage maps generated in this paper have been deposited to Zenodo (DOI: 10.5281/zenodo.17197823).

## Competing Interest Statement

The authors declare no competing interests.

## Acknowledgments

We thank Antoine Fraimout for access to data on laboratory crosses and Pasi Rastas for his generous advice in building the linkage maps. Our research was supported by grants from the Academy of Finland (# 129662, 134728 and 218343 to JM), grant from Helsinki Institute for Life Sciences (HiLife; to JM), a grant from the NSFC/RGC Joint Research Scheme sponsored by the Research Grants Council of the Hong Kong Special Administrative Region, China and the National Natural Science Foundation of China (Project No. N_HKU763/21) and two General Research Fund (GRF) 2024/25 grants from Research Grant Council, Hong Kong (GRF17126824 and GRF17104824). H.W., C.Z. and K.R. were supported by Faculty of Science (HKU) funding to J.M. We acknowledge CSC – IT Center for Science, Finland, for access to computational resources.

## Author Contributions

J.M., K.R., and C.Z. conceived and designed the study. H.W. constructed the linkage maps. C.Z. performed data analyses and plotting. C.Z., J.M., H.W., and K.R. wrote the paper.

## Notes

### Competing Interest Statement

The authors have declared no competing interest.

